# A GAL80 collection to inhibit GAL4 transgenes in *Drosophila* olfactory sensory neurons

**DOI:** 10.1101/168468

**Authors:** Jessica Eliason, Ali Afify, Christopher Potter, Ichiro Matsumura

**Affiliations:** Janelia Research Campus, Howard Hughes Medical Institute; Ashburn, VA, 20147; United States; Department of Biochemistry; Emory University School of Medicine; Atlanta, GA, 30322; United States; Solomon H. Snyder Department of Neuroscience; Johns Hopkins University School of Medicine; Baltimore, MD, 21205; United States

**Author notes:** 19700 Helix Drive; Ashburn, VA; 20147. Corresponding authors’ contact information Jessica Eliason 19700 Helix Drive 2E.120.2 Ashburn, VA 20147; & Ichiro Matsumura 1510 Clifton Rd NE 4119 Rollins Research Center Atlanta, GA 30322.

**Keywords:** Olfaction, GAL80, Drosophila, Odorant Receptor, Sensory Neurons

## Abstract

Fruit flies recognize hundreds of ecologically relevant odors and respond appropriately to them. The complexity, redundancy and interconnectedness of the olfactory machinery complicate efforts to pinpoint the functional contributions of any component neuron or receptor to behavior. Some contributions can only be elucidated in flies that carry multiple mutations and transgenes, but the production of such flies is currently labor-intensive and time-consuming. Here, we describe a set of transgenic flies that express the *Saccharomyces cerevisiae* GAL80 in specific olfactory sensory neurons (*OrX-GAL80s*). The GAL80s effectively and specifically subtract the activities of GAL4-driven transgenes that impart anatomical and physiological phenotypes. *OrX-GAL80s* can allow researchers to efficiently activate only one or a few types of functional neurons in an otherwise nonfunctional olfactory background. Such experiments will improve our understanding of the mechanistic connections between odorant inputs and behavioral outputs at the resolution of only a few functional neurons.

## INTRODUCTION

The olfactory system of *Drosophila melanogaster* is often the subject in studies of memory, evolution, gene choice, development and odorant-induced behavior. It is a good model system because of its relatively stereotyped neuronal circuitry, complex behaviors and convenient genetic tools.

In *Drosophila*, most olfactory sensory neurons (OSNs) typically expresses a single odorant receptor (OR) from a genomic repertoire of 60 genes (Vosshall et al. 1999; Robertson, Warr, and Carlson 2003; Vosshall, Wong, and Axel 2000; Clyne et al. 1999; Goldman et al. 2005). The promoter of an OR gene can be employed to label specific subsets of OSNs with a particular transgene (Fishilevich and Vosshall 2005; Couto, Alenius, and Dickson 2005). ORs, which vary in sensitivity and specificity to a wide range of different odorants, determine the firing kinetics and odor-response dynamics of each OSN (Hallem and Carlson 2006; Hallem, Ho, and Carlson 2004; Couto, Alenius, and Dickson 2005; Fishilevich and Vosshall 2005; de Bruyne, Clyne, and Carlson 1999; de Bruyne, Foster, and Carlson 2001; Dobritsa et al. 2003; Elmore et al. 2003; Kreher et al. 2008).

Most OSNs express Odorant Receptor Co-Receptor (Orco), a highly conserved member of the olfactory receptor family (Krieger et al. 2003; Vosshall and Hansson 2011), in addition to a single selected OR. Though Orco usually does not contribute to the structure of the odorant binding site (Jung, borst, and Haag 2011; Nakagawa and Vosshall 2009; Nichols and Luetje 2010; P. L. Jones, Pask, and Rinker 2011), it is essential for odorant-invoked signaling in flies. Without Orco, the co-expressed OR cannot localize to the dendritic membrane or relay an odor-evoked signal (Larsson et al. 2004; Benton et al. 2006). Orco null flies are largely anosmic, though some chemosensation remains due to the presence of ionotropic receptors and gustatory receptors, which do not require Orco to function (Silbering et al. 2011; Benton et al. 2009; Kwon et al. 2007; W. D. Jones et al. 2007). The Orco promoter is consequently a convenient device for the expression of transgenes in most OSNs.

The olfactory organs, the antenna and maxillary palp, contain OSNs dendrites within structures called sensilla. ORs and Orco are embedded in the dendritic membrane. OSN axons project to the antennal lobes in the brain of the animal. Each antennal lobe consists of ~50 globular synaptic sites called glomeruli. All OSNs on the periphery that expresses the same OR converge onto their own unique glomerulus. For example, all OSNs expressing Or22a will send axons to the DM2 glomerulus in the antennal lobe while all OSNs expressing Or82a will send axons to the VA6 glomerulus (**Figure 1**). The stereotyped organization of OSNs and their projections is known as the olfactory sensory map (Vosshall, Wong, and Axel 2000; Fishilevich and Vosshall 2005; Couto, Alenius, and Dickson 2005; Stocker et al. 1990). The regularity of this map is a key feature that makes *Drosophila* olfaction such a useful model, as any aberration to the typical pattern will be apparent. The apparent simplicity of the map (**Figure 1**), however, obscures mechanistic complexities that are yet to be discovered, in part because necessary tools remain unavailable.

**Figure 1:**
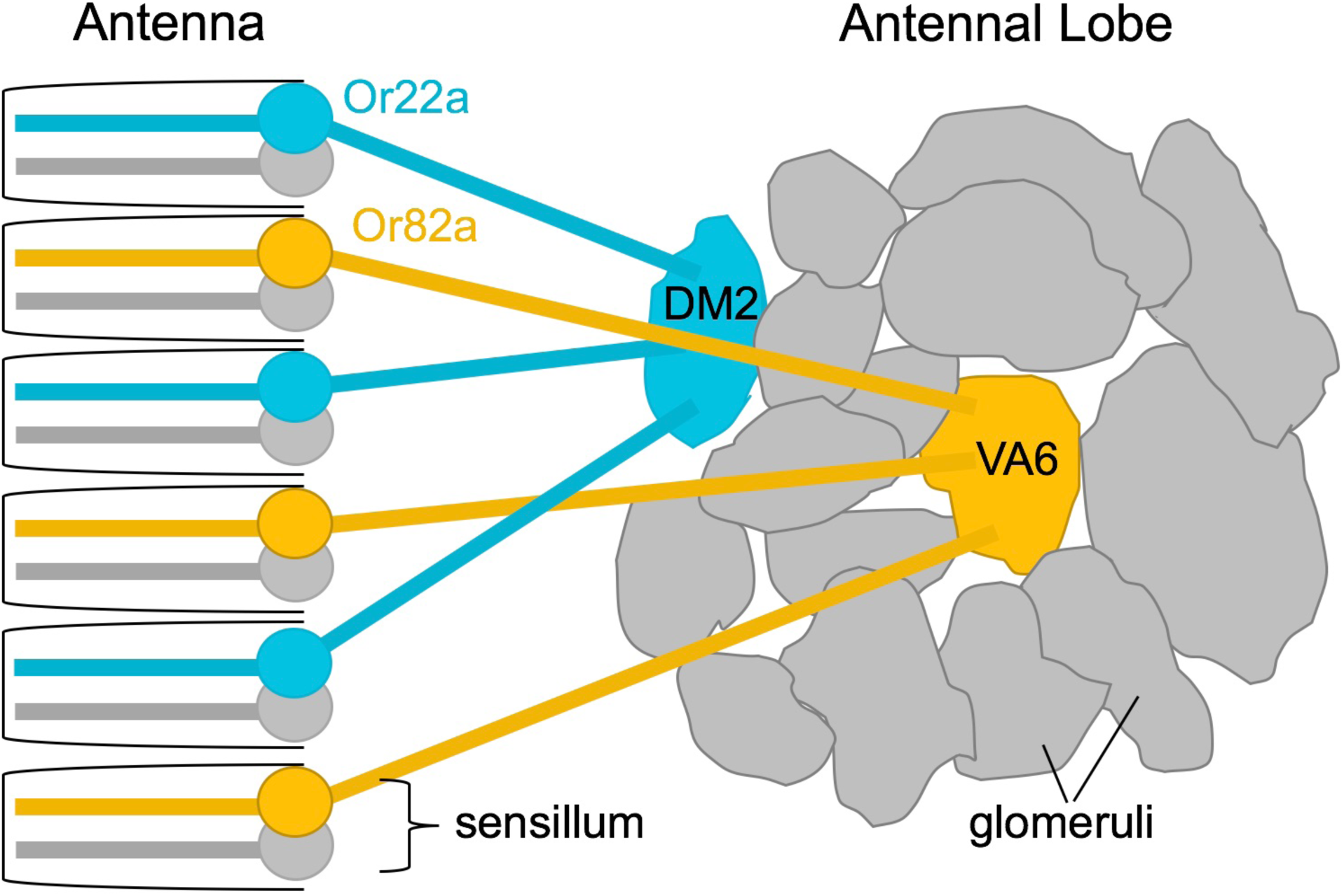
Olfactory Sensory Map. Each neuron in the olfactory system expresses one type of odorant receptor (OR). Or22a (teal) and Or82a (gold) are used here as examples. Neurons usually exist in pairs or groups in sensilla within the olfactory organs—antenna or maxillary palp. Neurons expressing the same OR are distributed throughout the periphery, but project their axons onto the same glomerulus in the antennal lobe of the brain. For example, all Or22a-expressing neurons synapse onto the DM2 glomerulus while all Or82a-expressing neurons synapse onto the VA6 glomerulus.

*Drosophila* geneticists have traditionally relied on genetic mutations or deletions to understand how complex biological systems normally work. Most alleles are recessive, so homozygotes must be bred over multiple generations. Achieving homozygosity of a mutation while also adding transgenes to the system often requires the creation of recombinant chromosomes produced after multiple generations of crossing and PCR screening. Classical genetic strategies thus limit the number and complexity of combinatorial genotypes that one can achieve. More challenging experimental questions demand more facile and versatile genetic tools.

The GAL4/UAS gene regulation system has become a *defacto* standard in studies of *Drosophila*. GAL4 is a yeast transcription activator that binds to the Upstream Activating Sequence (UAS) and induces expression of downstream genes (Giniger, Varnum, and Ptashne 1985). By driving GAL4 expression from an OR promoter, specific expression of a *UAS-transgene* can be obtained for any OSN subtype. An *OrX-GAL4* line exists for almost every OR. This collection of GAL4 lines is a powerful toolbox since different *UAS-transgenes* can be introduced into a line via conventional mating. For example, human α-synuclein has been expressed in OSNs to model human Parkinson’s disease (A. Y. Chen et al. 2014). Alternatively, protein expression levels can be knocked down using any specified *UAS-RNAi* transgene.

A variety of existing compatible effectors can be used study different aspects of neuronal communication. The *UAS-Kir2.1* effector is used as an example in experiments described below. This inward rectifier potassium channel electrically inactivates the neurons that express it (Hodge 2009; Baines et al. 2001; Johns et al. 1999). Similarly, *shibire^ts^* or tetanus toxin can be used to silence synaptic communications (van der Bliek and Meyerowitz 1991; kitamoto 2002; Kitamoto 2001; M. S. Chen et al. 1991; Sweeney et al. 1995; Baines et al. 1999), *reaper/grim/hid* genes can be used to physically kill neurons by inducing their own apoptotic pathways (Song and Steller 1999; Abrams 1999), or ricin toxin can be expressed ectopically to kill neurons. Conversely, neurons can be selectively activated with *trp1a* or a variety of other channelrhodopsin transgenes (Boyden 2011; Pulver et al. 2009).

If GAL4 is a standard on-switch for nearly any desired transgene, GAL80 is the logical off-switch. GAL80 binds the GAL4 transcriptional activation domain, thereby preventing recruitment of RNA polymerase (Ma and Ptashne 1987). GAL80 crosses are much more convenient than classical breeding approaches (**Figure 2**). In order to have a single functional OSN in an otherwise silent olfactory system, the traditional method uses an Orco null mutation (Larsson et al. 2004). In this genetic setup, Orco mutant flies are mostly anosmic, but function is restored to one OSN subset with *Or-GAL4, UAS-Orco* transgenes (Olsen, Bhandawat, and Wilson 2007; DasGupta and Waddell 2008; Hoare, McCrohan, and Cobb 2008; Hoare et al. 2011; Benton et al. 2006; Fishilevich et al. 2005) (**Figure 2a**). An *Orco-GAL4, UAS-effector, Or-GAL80* method can be used instead (**Figure 2b**). Kir2.1 is used as an example of an effector (Hodge 2009; Baines et al. 2001; Johns et al. 1999). Classical breeding strategies (**Figure 2a**) may look less complicated on paper than GAL80 crosses (**Figure 2b**) but are actually more time-consuming and limited. The Orco mutation must be homozygous. Since most *Drosophila* transgenes are embedded into the same two chromosomes (2 or 3) recombination and PCR screening may be required to achieve this homozygosity.

**Figure 2:**
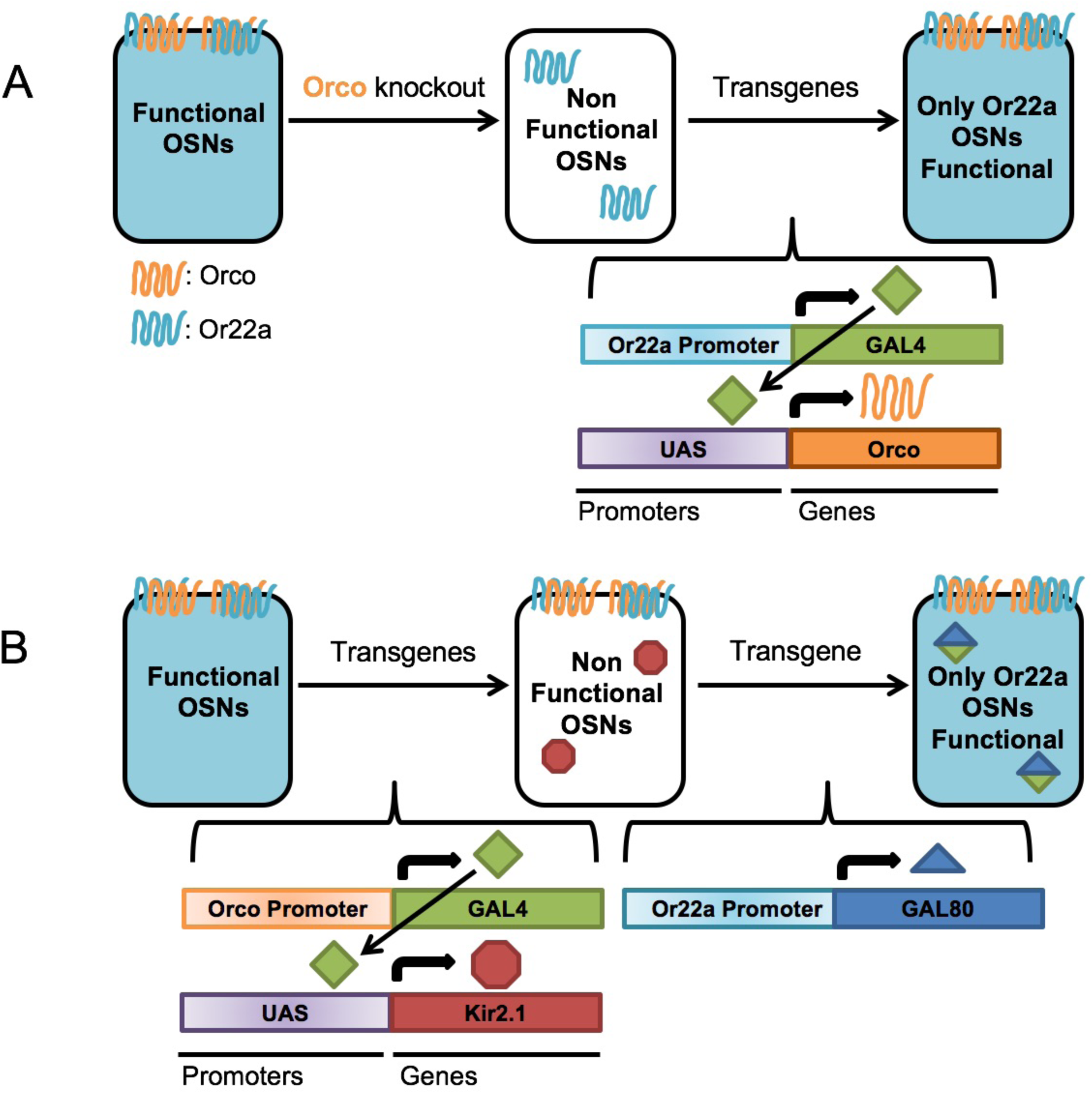
Advantages of using a GAL80 approach over a null mutation. a) **Current method with available reagents**. In order to examine a single type of Olfactory Sensory Neuron (OSN) without interference from other OSNs, one can use an Orco null mutant. Without Orco, ORs cannot reach the cell membrane or function properly. Orco mutants are mostly anosmic (unable to smell.) A single OR can then be restored using two transgenes, *OrX-GAL4* and *UAS-Orco. Or22a-GAL4* is shown here as an example. This fly may require the making and validating of one or more recombinant chromosomes, since the Orco mutation must be homozygous. In more complicated systems, e.g. restoring more than one OSN, multiple recombinants would need to be made and validated at a cost of several months of crossing. b) **Using a GAL80**. GAL80 is a potent GAL4 inhibitor. All olfactory neurons could be silenced using any number of transgenes in an *Orco-GAL4, UAS-effector* (such as *UAS-Kir2.1*) genotype. A single OSN subtype can then be restored using an *OrX-GAL80* (such as *Or22a-GAL80*). This system requires no recombinant creation, and is amenable to the use of various effectors or additional transgenes without requiring recombinant construction. (Receptor appearance, orientation, and heterodimerization is based on previous designs by Neuhaus et al. (2005), Benton et al. (2006), Smart et al. (2008), and (Smart et al. 2008; Neuhaus et al. 2005; Benton et al. 2006; Benton 2009)

Neurons seldom operate autonomously, but rather groups of neurons coordinate within a circuit to provide an organism with perception and behavior. An investigation of the behavioral impact provided by a limited number of different functional neuronal types would require additional genes. The elaboration of genotypes (**Figure 2**) to restore pairs or groups of functional OSNs in a nonfunctional background normally requires generations of crosses (followed by PCR screens for desired recombinants). A GAL80 strategy can shorten this process by achieving similar results in only one or two generations with no necessary recombinant creation. Furthermore, a GAL80 strategy takes advantage of the interchangeable variety of existing *UAS-transgene* lines. Here we describe a new collection of *OrX-GAL80* lines designed to complement existing *OrX-GAL4* lines, and demonstrate their potential utility for neuroanatomical studies of the *Drosophila* olfaction model.

## MATERIALS AND METHODS

### Fly Stocks

Flies were reared on standard cornmeal/molasses food and kept at 25C with a 16 hours on/8hours off light cycle. All lines were obtained from the Indiana University Bloomington Stock Center and the Janelia Research Campus. Any recombinants made were validated with PCR.

Stock List
Or7a-GAL4 #23907 Or7a-GAL4 #23908 Or10a-GAL4 #9944 Or13a-GAL4 #9946 Or13a-GAL4 #23886 Or19a-Gal4 #24617 Or22a-GAL4 #9951 Or22a-GAL4 #9952 Or22b-GAL4 #23891 Or33c-GAL4 #23893 Or35a-GAL4 #9967 Or42a-GAL4 #9970 Or42b-GAL4 #9971 Or43b-Gal4 #23894 Or46a-GAL4 #23291 Or47a-GAL4 #9981 Or56a-GAL4 #9988 Or59b-GAL4 #23897 Or59c-GAL4 #23899 Or67a-GAL4 #23904 Or67d-GAL4 #9998 Or71a-GAL4 #23121 Or82a-GAL4 #23125 Orco-GAL4 #23292 Orco-GAL4 #26818 Or85a-GAL4 #23133 Or85b-GAL4 #23911 Or85c-GAL4 #23913 Gr21a-GAL4 #24147 Or22a-mcd8::GFP #52620 Gr21a-mcd8::GFP #52619 Orco^2^ #23130 UAS-Orco #23145 UAS-mcd8::GFP #5130 UAS-mcd8::GFP #5137 UAS-Kir2.1 Janelia stock #3015545 UAS-Kir2.1 Janelia stock #3015298 UAS-Kir2.1::eGFP Janelia stock #BS00312 pJFRC19-13xLexAop2-IVS-myr::GFP-p10 (attP8) Janelia stock #1171146 pJFRC59-13xLexAop2-IVS-myr::GFP-p10 (attP40) Janelia stock #3015445

### GAL80 Creation

Primers were designed to capture the entire promoters described by (Couto, Alenius, and Dickson 2005) (see **Table S1**). Promoters were amplified from genomic DNA using Q5 High Fidelity PCR (NEB #M0491S) and added to entry vectors using the pENTR/D-TOPO system (Invitrogen 2012b). Recombination with the pBP-GAL80Uw-6 (Addgene #26236) destination vector was done using the LR Clonase II system (Invitrogen 2012a). To ensure no mutations, no gaps, and correct orientation, the complete promoters were sequenced in the destination vector using the sequencing primers shown in **Table S2**. PhiC31 site-directed transgenesis was performed by Genetivision Inc. All GAL80 transgenes were inserted at the attP2 site. A single Or-LexA line was also created using the Or22a-promoter entry vector and pBPnlsLexA::p65Uw (Addgene #26230).

### Immunohistochemistry

Female adult brains were dissected one day after eclosion in cold S2 Schneider’s Insect Medium (Sigma Aldrich #S0146) and fixed while nutating for 55 minutes at room temperature in 2mL 2%PFA (Electron Microscopy Sciences #15713) in protein loBind Tubes (Eppendorf #022431102). Brains were washed 4x, 15min per wash while nutating with 2mL PBT buffer (1xPBS, Cellgro #21-040, with 0.5% TritonX-100, Sigma Aldrich #X100). Brains were then blocked with 200μL 5% Goat serum (ThermoFischer. #16210064) in PBT for 90 minutes while nutating, upright. Block was removed and 200 μL primary antibodies in PBT were added for 4 hours at room temperature and then transferred to 4C for 36-48 hours while nutating, upright. Primary antibodies: mouse α-bruchpilot (Developmental Studies Hybridoma Bank. #nc82-s) at 1:30, rabbit α-GFP at 1:1000 (Thermo Fischer #A11122), or rabbit α-Tom at 1:500 (clontech #632393). Monoclonal antibody nc82 identifies Bruchpilot. Bruchpilot can serve as a general neuropil marker because it is required in synaptic zones (Wagh et al. 2006). Larval brains were collected from third instar larvae and fixed in 4% PFA. Primary antibodies: mouse α-neuroglian (Developmental Studies Hybridoma Bank. #BP104) at 1:50 and rabbit α-GFP at 1:500. Brains were washed 4x, 15min per wash while nutating with 2mL PBT. 200μL secondary antibodies in PBT were then added for 4 hours at room temperature and then 3 overnights at 4C while nutating upright. Secondary antibodies: AF568 goat α-mouse (Life Technologies #A11031) at 1:400 and AF488 goat α-rabbit (ThermoFischer #A11034) at 1:800. Tubes were protected from light at all times after secondary antibodies had been added. Brains were washed again 4x, 15min per wash while nutating with 2mL PBT. Then washed with 1xPBS and mounted using Vectashield mounting media (Vector Labs #H-1000). Confocal images were taken with Leica800 microscope.

### Single Sensillum Recordings

SSRs were performed as described in Lin et al 2015 (Lin and Potter 2015). GFP labeled ab1 and ab3 sensilla were identified using a Zeiss AxioExaminer D1 compound microscope with eGFP filter cube (FL Filter Set 38 HE GFP shift free). A glass recording electrode filled with ringers solution (7.5g of NaCl+0.35g of KCl+0.279g of CaCl_2_-2H_2_O in 1L of H_2_O) was inserted into the base of the sensillum. To test ab1 (Gr21a) response, CO_2_ was delivered through a tube ending with a Pasteur pipette that was inserted for 1 second into a hole in a plastic pipette directed at the antenna. This plastic pipette (Denville Scientific Inc, 10ml pipette) carried a purified continuous air stream (8.3 ml/s) that used a stimulus controller (Syntech) at the time of CO_2_ delivery to correct for the increased air flow. To test ab3 (Or22a) response, 20 μl of E2-Hexenal or Isoamyl acetate (diluted to 1% in mineral oil) was pipetted on a piece of filter paper (1X2 cm) in a Pasteur pipette. The Pasteur pipette was then inserted into the hole of the plastic pipette that carried continuous air stream to the antenna. For odorant delivery, the stimulus controller (Syntech) was used to divert a 1 s pulse of charcoal-filtered air (5 ml/s) into the Pasteur pipette containing the odorant.

Signals were acquired and analyzed using AUTOSPIKE software (USB-IDAC System; Syntech). Spikes were counted in a 500 ms window from 500 ms after CO_2_ delivery and multiplied by 2 to calculate spikes/second. Then, the spikes in 1000ms before CO_2_ delivery were subtracted to calculate the increase in spike rate in response to CO_2_ (Δspikes/second). For each genotype, 6 flies (4-8 days old) were tested, with 1-3 sensilla tested in each fly.

## RESULTS

### Design of GAL80 Constructs

The following criteria were used to choose OR promoters for the collection. i) The ORs should be relevant to current research as shown by the number of studies that used it. ii) The ORs should represent a variety of expression patterns (larval or adult, antennae or maxillary palps, sensillary class etc.). iii) Finally, the ORs should reflect a variety of different odorant response profiles. The promoter regions were defined based largely on the work of Couto *et al*, 2005.

Equimolar expression of GAL4 and GAL80 is not always sufficient to effectively eliminate GAL4 activity so the pBP-GAL80uW-6 vector was used. This vector contains a modified GAL80 sequence, designed to increase the stability and expression of its gene product (Pfeiffer et al. 2010). A few *OrX-GAL80s* were already made with this vector and used effectively. (Gao, Clandinin, and Luo 2015) pioneered the technique by creating a limited number of OrX-GAL80s. This work is a logical extension and makes many additional *OrX-GAL80s* available for general use.

### Testing GAL80 Efficacy and Specificity

GAL80 lines were created for the following odorant receptor promoters: Or7a, Or10a, Or13a, Or19a, Or22a, Or22b, Or33c, Or35a, Or42a, Or42b, Or43b, Or47a, Or56a, Or59b, Or59c, Or67a, Or67d, Or71a, Or82a, Orco, Or85a, Or85b, Or85c, and Gr21a. To examine GAL4 subtraction *in vivo, OrX-GAL80* flies were crossed to flies with the genotype *OrX-GAL4, UAS-GFP*. OSNs expressing the same OR can be identified from their specific glomerulus in the antennal lobe (**Figure 1**). *OrX-GAL4, UAS-GFP* flies show robust expression of the GFP reporter gene in their respective glomeruli. However, when *OrX-GAL80* is added to the genotype, GFP expression is entirely absent, indicating a robust antagonism of GAL4 activity (**Figure 3**). The efficacy of *Or22b-GAL80* could not be determined because the *OR22b-GAL4, UAS-GFP* control did not show robust or reliable GFP signaling in the first place. The created *Or7a-GAL80* line was not effective at subtracting GFP signal. Though these lines are not included in Figure 3, they will still be available in the Bloomington Stock Center. Several of the GAL80 lines also have expression in larvae. GAL4 subtraction was examined in larval brains using the *UAS-GFP* reporter gene. In larvae, GAL80 reduced but did not eliminate GAL4 activity (**Figure S1a**).

**Figure 3:**
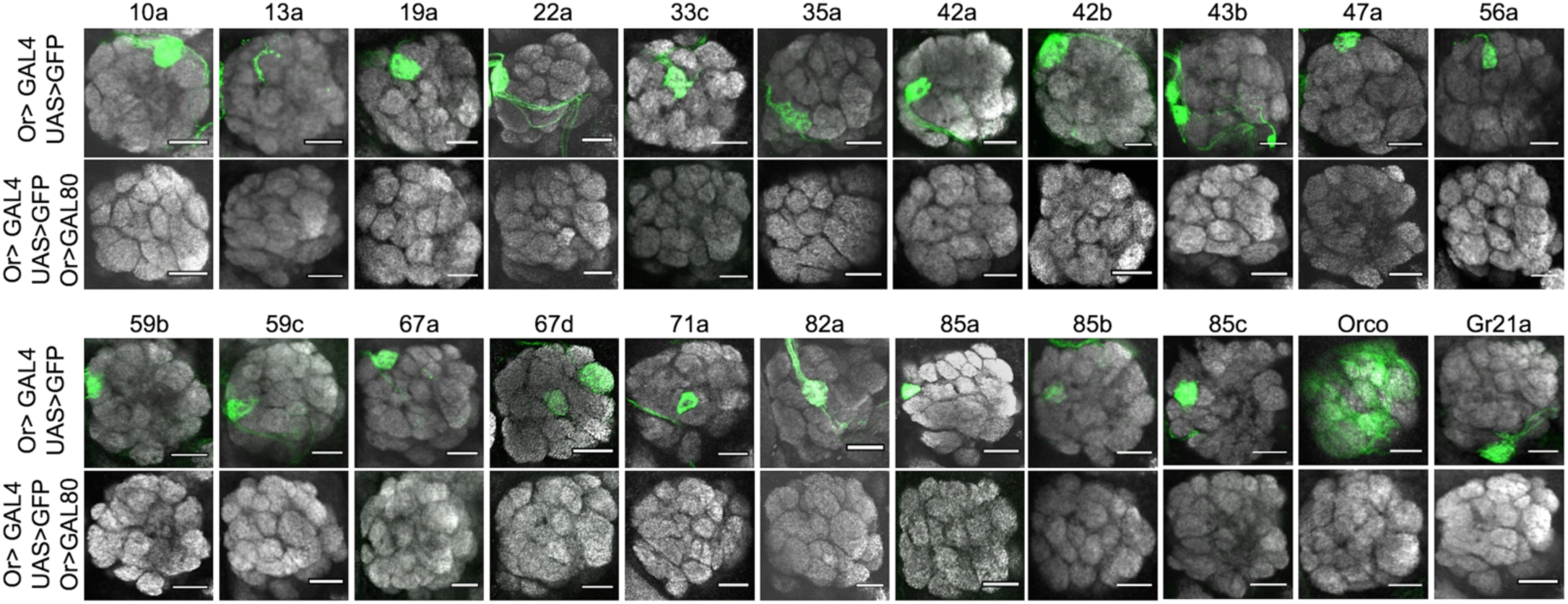
OR-GAL80 reagents eliminate GAL4 activity. All antennal lobes are stained with anti-nc82 (a general neuropil marker, grey) and anti-GFP (green). The orientation of each image is dorsal-up, ventral-down, lateral-right, medial-left. Scale bars indicate 20μm. Each of the brains shown has the genotype *OrX-GAL4, UAS-GFP*. The specific receptor promoter is given above each column. The top row in each set shows GFP expression in these lines without GAL80. Notice how each neuron’s target in the antennal lobe glomeruli is expressing GFP. Each bottom row shows the brains containing an additional *Or-GAL80* gene. Note how GAL80 effectively inhibits GAL4 activity, as seen by the elimination of GFP expression. The images are representative of the 5-20 brains examined per genotype. GAL4 inactivation was 100% penetrant in one day old female flies.

The *OrX-GAL80* lines were checked to ensure they would not have aberrant expression in untargeted OSN subtypes. The pBP-GAL80uW-6 vector contains a Drosophila Synthetic Core Promoter (DSCP). DSCP is an effective means of using enhancer elements to drive strong expression (Pfeiffer et al. 2008), but it could also cause the GAL80s to have nonspecific or leaky expression. Therefore, a version of pBP-GAL80uW-6 was cloned with the DSCP removed. However, when the DSCP was absent, GAL80 expression was insufficient to subtract GAL4 activity (**Figure S1b**). A few lines were tested to see if DSCP causes nonspecific GAL80 expression. For these lines, an *OrY-GAL80* did not impede GAL4 activity of an *OrX-GAL4* neuron (**Figure S1c**). Due to the uneven expression in an *Orco-GAL4, UAS-GFP* line, it could not be determined if each OrX-GAL80 subtracts GAL4 from only one glomerulus in an otherwise fully-labeled brain, but results shown in Figure S1c give reasonable confidence that the GAL80s do not have widespread nonspecific expression. It can also be noted that the GAL80 subtraction does not interfere with reporter gene expression in a genetic system that does not use GAL4. When *Or22a-GAL80* is used in conjunction with *Or22a-GFP*, containing no GAL4/UAS intermediary, the GFP is still expressed (**Figure S1d**). These images, showing subtraction of reporter gene expression, confirm that GAL4 activity is suppressed anatomically by the GAL80 lines.

To confirm GAL4 was suppressed physiologically by the GAL80s, Single Sensillum Recordings (SSRs) were used to assay electrical activity of OSNs. *Gr21a-GFP* was used to identify sensilla of interest without interfering with the GAL4/UAS/GAL80 system. Gr21a neurons are housed in ab1 sensilla. Carbon Dioxide exposure causes a robust response in Gr21a ab1C neurons (Hallem and Carlson 2006; Hallem, Ho, and Carlson 2004; Fishilevich and Vosshall 2005; deBruyne:2001bs W. D. Jones et al. 2007; Kwon et al. 2007; de Bruyne, Foster, and Carlson 2001). When *Gr21a-GFP* flies were exposed to CO_2_, their ab1C sensillar neurons showed robust responses (mean Δspikes/s=88, N=8 sensilla). Kir2.1-containing neurons are expected to show little to no spontaneous firing (Olsen, Bhandawat, and Wilson 2007; Hoare, McCrohan, and Cobb 2008). Adding Kir2.1 to Gr21a neurons (genotype *Gr21a-GFP, Gr21a-GAL4, UAS-Kir2.1*) greatly reduced spiking responses to CO_2_ (mean Δspikes/s=14, N=12 sensilla, p=0.01). When Gr21a-GAL80 was added (genotype *Gr21a-GFP, Gr21a-GAL4, UAS-Kir2.1, Gr21a-GAL80*), responses to CO_2_ were restored (mean Δspikes/s=94, N=6 sensilla, *p*<0.001. No significant difference from genotype *Gr21a-GFP*, p= .26) (**Figure 4a**).

**Figure 4:**
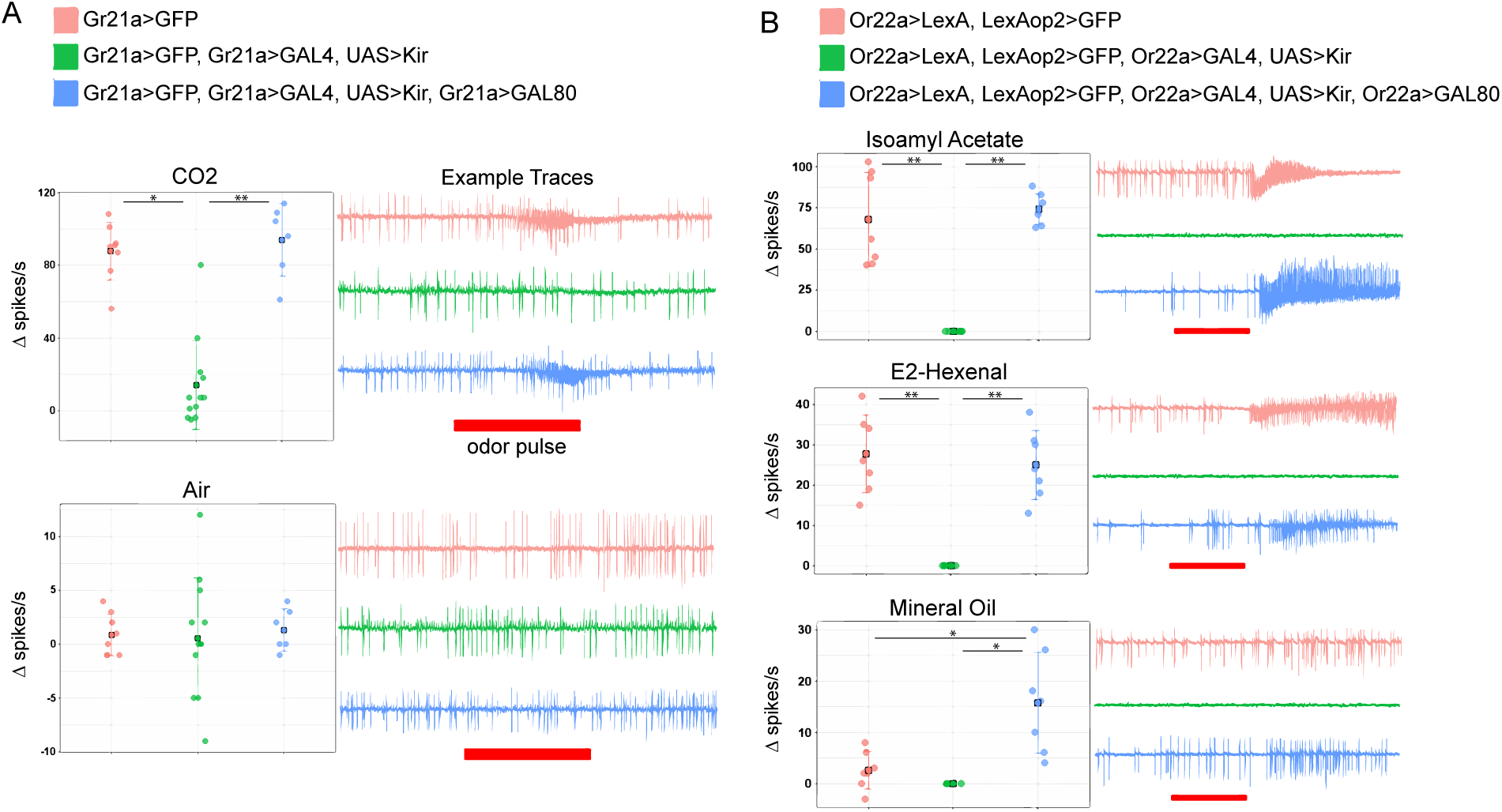
Olfactory neuron responses towards odors in Single Sensillum Recordings (SSR). In box plots on the left, each circle shows response in an individual sensillum, and filled squares indicate the means. On the right of each plot, example SSR traces are shown for each genotype. (* indicates .01>p>.005, ** indicates p<.001) a) **Ab1C SSR responses**. Ab1C neurons are visualized using the *Gr21a-GFP* gene. Top: Sensilla respond strongly to CO_2_, and adding *Kir2.1* reduces response to CO_2_. Response is restored when *Gr21a-GAL80* is added. Bottom: Air was used as a control for CO_2_ experiments. Air does not cause an odor-evoked neuronal response, and adding the *Kir2.1* or *GAL80* genes does not affect the spontaneous signaling responses. b) **Ab3 SSR responses**. Ab3 neurons are visualized using the *Or22a-LexA and LexAop2-GFP* genes. Sensilla respond to Isoamyl Acetate and to E2-Hexenal. Adding *Kir2.1* eliminates both odor-evoked and spontaneous activity in these neurons. Spontaneous and odor-evoked activity is restored when *Or22a-GAL80* is added. Odorants were diluted in mineral oil and neurons from the GAL80 restorative genotype did show low-level responses to mineral oil alone (bottom).

To make sure the system also worked for neurons expressing an OR protein (in additional to a GR), SSR was also done with ab3 sensilla. Ab3 houses Or22a-expressing neurons. This receptor is known to be activated by a diverse set of odorants, including Isoamyl acetate and E2-hexenal (Hallem and Carlson 2006; Hallem, Ho, and Carlson 2004; Fishilevich and Vosshall 2005; deBruyne:2001bs W. D. Jones et al. 2007; Kwon et al. 2007; de Bruyne, Foster, and Carlson 2001). A transgene was necessary to visualize the neurons without interfering with the GAL4/UAS/GAL80 system, but *Or22a-GFP* was insufficiently bright to identify sensilla for SSR. A new *Or22a-LexA* transgenic animal was therefore created using the same promoter that was used to create the *Or22a-GAL80* gene (this line is also available through Bloomington). When crossed to a *LexAop2-GFP* line, the ab3 sensilla showed bright fluorescence. When *Or22a-LexA, LexAop2-GFP* flies were exposed to Isoamyl acetate or to E2-hexenal, their sensillar neurons showed robust responses (mean Δspikes/s=67.86 and 27.71, N=7 and 7 sensilla, respectively). Unlike the Gr21a neurons, Kir2.1 expression in the Or22a neurons effectively eliminated both spontaneous and odor-evoked activity. (mean Δspikes/s=0, N=7 sensilla, p<.001 for both odorants). Both activities could be restored with the addition of the *Or22a-GAL80* gene (Isoamyl acetate: mean Δspikes/s =74.29, N=7 sensilla, p<.001; E2-Hexenal: mean Δspikes/s =25, N=7 sensilla, p<.001). Neurons showed some low-level responses to mineral oil alone, the solvent used for the odorants (**Figure 4b**). Only genotype 3 *Or22a-LexA, LexAop2-GFP, Or22a-GAL4, UAS-Kir2.1, Or22a-GAL80* showed significantly higher responses to mineral oil than genotypes *Or22a-LexA, LexAop2-GFP* (p=.01) and *Or22a-LexA, LexAop2-GFP, Or22a-GAL4, UAS-Kir2.1* (p=.005), but the latter two genotypes showed no significant response to mineral oil alone. The results in **Figure 4** confirm that GAL80 functions effectively to prevent GAL4-induced activity in OSNs.

## DISCUSSION

The collection of GAL80 lines subtracts GAL4 activity efficiently and specifically in OSNs. In anatomical studies, reporter gene expression from the GAL4/UAS system is suppressed. Neurons silenced with Kir2.1 expression have normal firing capacity restored when GAL4 is antagonized using the GAL80 lines.

In behavioral assays, using a GAL80 transgene will be more flexible than mutant lines and less cumbersome than crafting the required recombinants as the complexity of the genotype increases. Though in some special circumstances, olfactory sensory neurons have been shown to produce behaviors autonomously, this is not a widely applicable principle, and further investigation upon this principle requires better tools. For example, Fishilevish et al (2005) used larvae in their study to restore aversion with a single functional OSN subtype, but the larval olfactory system may be fundamentally different from adults in this respect., (Bhandawat et al. 2010) also showed that single glomerular activity is sufficient to invoke a behavioral response, but that study was done using an intact and fully functional olfactory background, so some neuronal cooperation may still have occurred. (DasGupta and Waddell 2008) provided evidence that a single functional OSN subtype is sufficient to learn odor discrimination, and Gao et al (2015) gave convincing evidence of aversive restoration in adults with only one functional OSN.

However, the extent to which the restoration of single-OSN behavior depends on the odorant and receptor used is still unknown. Only a small subset of receptors have been tested. The current models of odor coding by the olfactory system predict that a coordinated effort of many OSNs is usually required to produce a behavioral output. Paired neurons in sensilla can affect the firing dynamics of their neighbors in the periphery (Dobritsa et al. 2003; Su et al. 2012; Kazama and Wilson 2009), and downstream neurons such as interneurons and projection neurons may rely on synchronized input from multiple OSN types (Chou et al. 2010; Wilson 2011; Yaksi and Wilson 2010; Hong and Wilson 2015; Kazama, Yaksi, and Wilson 2011; Olsen, Bhandawat, and Wilson 2007; Ng et al. 2002; Acebes et al. 2011). GAL80 tools open more possibilities to combinatorially activate subsets of neurons. The hope is that additional researchers will use the reagents and validate them in their own assays.

Researchers encounter a significant technical obstacle to the understanding of olfactory function if they need to create genotypes with small groups of interacting neurons in isolation. The tools presented here facilitate the activation or deactivation of combinations of particular neurons, thereby overcoming this obstacle. The lines are available to order through Bloomington Stock Center.

## AUTHOR CONTRIBUTIONS

Conceptualization, JE and IM; Methodology, JE; Formal Analysis, JE and AA; Investigation, JE and AA; Writing—Original Draft, JE; Writing—Review and Editing, AA, CP, IM; Visualization, JE and AA; Supervision, IM; Funding Acquisition, IM

## ACKNOWLEDGEMENTS

Funding for this project comes from the National Science Foundation (MCB-1413062). The authors report no conflicts of interest. We thank Heather Dionne for cloning advice, Yoshi Aso for the use of his behavioral setup and Janelia FlyLight for consulting on imaging and immunostaining.

## REFERENCES

Abrams, J M. 1999. “An Emerging Blueprint for Apoptosis in Drosophila.” Trends in Cell Biology 9 (11): 435–40.

Acebes, Angel, Alfonso Martín-Peña, Valérie Chevalier, and Alberto Ferrús. 2011. “Synapse Loss in Olfactory Local Interneurons Modifies Perception.” The Journal of Neuroscience : the Official Journal of the Society for Neuroscience 31 (8): 2734–45. doi:10.1523/JNEUROSCI.5046-10.2011.

Baines, R A, S G Robinson, M Fujioka, J B Jaynes, and M Bate. 1999. “Postsynaptic Expression of Tetanus Toxin Light Chain Blocks Synaptogenesis in Drosophila.” Current Biology 9 (21): 1267–70.

Baines, Richard A, Jay P Uhler, Annemarie Thompson, Sean T Sweeney, and Michael Bate. 2001. “Altered Electrical Properties in DrosophilaNeurons Developing Without Synaptic Transmission.” Journal of Neuroscience 21 (5). Society for Neuroscience: 1523–31. doi:10.1016/S0960-9822(99)80510-7.

Benton, R. 2009. “Evolution and Revolution in Odor Detection.” Science 326 (5951): 382–83. doi:10.1126/science.1181998.

Benton, Richard, Kirsten S Vannice, Carolina Gomez-Diaz, and Leslie B Vosshall. 2009. “Variant lonotropic Glutamate Receptors as Chemosensory Receptors in Drosophila.” Cell 136 (1): 149–62. doi:10.1016/j.cell.2008.12.001.

Benton, Richard, Silke Sachse, Stephen W Michnick, and Leslie B Vosshall. 2006. “Atypical Membrane Topology and Heteromeric Function of Drosophila Odorant Receptors in Vivo.” PLoS Biology 4 (2): e20–18. doi:10.1371/journal.pbio.0040020.

Bhandawat, V, G Maimon, M H Dickinson, and R I Wilson. 2010. “Olfactory Modulation of Flight in Drosophila Is Sensitive, Selective and Rapid.” Journal of Experimental Biology 213 (21): 3625–35. doi:10.1242/jeb.040402.

Boyden, Edward S. 2011. “A History of Optogenetics: the Development of Tools for Controlling Brain Circuits with Light.” F1000 Biology Reports 3 (11): 11. doi:10.3410/B3-11.

Chen, Alex Y, Shouzhen Xia, Paul Wilburn, and Tim Tully. 2014. “Olfactory Deficits in an Alpha-Synuclein Fly Model of Parkinson’s Disease.” Edited by Mel B Feany. PLoS ONE 9 (5): e97758. doi:10.1371/journal.pone.0097758.

Chen, M S, R A Obar, C C Schroeder, T W Austin, C A Poodry, S C Wadsworth, and R B Vallee. 1991. “Multiple Forms of Dynamin Are Encoded by Shibire, a Drosophila Gene Involved in Endocytosis.” Nature 351 (6327): 583–86. doi:10.1038/351583a0.

Chou, Ya-Hui, Maria L Spletter, Emre Yaksi, Jonathan C S Leong, Rachel I Wilson, and Liqun Luo. 2010. “Diversity and Wiring Variability of Olfactory Local Interneurons in the Drosophila Antennal Lobe.” Nature Neuroscience, February. Nature Publishing Group, 1–13. doi:10.1038/nn.2489.

Clyne, P J, C G Warr, M R Freeman, D Lessing, J Kim, and J R Carlson. 1999. “A Novel Family of Divergent Seven-Transmembrane Proteins: Candidate Odorant Receptors in Drosophila.” Neuron 22 (2): 327–38.

Couto, Africa, Mattias Alenius, and Barry J Dickson. 2005. “Molecular, Anatomical, and Functional Organization of the Drosophila Olfactory System.” Current Biology 15 (17): 1535–47. doi:10.1016/j.cub.2005.07.034.

DasGupta, Shamik, and Scott Waddell. 2008. “Learned Odor Discrimination in Drosophila Without Combinatorial Odor Maps in the Antennal Lobe.” Current Biology 18 (21): 1668–74. doi:10.1016/j.cub.2008.08.071.

de Bruyne, M, P J Clyne, and J R Carlson. 1999. “Odor Coding in a Model Olfactory Organ: the Drosophila Maxillary Palp.” The Journal of Neuroscience : the Official Journal of the Society for Neuroscience 19 (11). Society for Neuroscience: 4520–32.

de Bruyne, Marien, Kara Foster, and John R Carlson. 2001. “Odor Coding in the Drosophila Antenna.” Neuron 30 (2): 537–52. doi:10.1016/S0896-6273(01)00289-6.

Dobritsa, Anna A, Wynand van der Goes van Naters, Coral G Warr, R Alexander Steinbrecht, and John R Carlson. 2003. “Integrating the Molecular and Cellular Basis of Odor Coding in the Drosophila Antenna.” Neuron 37 (5): 827–41.

Elmore, Tamara, R Ignell, John R Carlson, and Dean P Smith. 2003. “Targeted Mutation of a Drosophila Odor Receptor Defines Receptor Requirement in a Novel Class of Sensillum | Journal of Neuroscience.” The Journal of Neuroscience : the Official Journal of the Society for Neuroscience 23 (30). Society for Neuroscience: 9906–12. doi:10.1038/81774.

Fishilevich, Elane, Ana I Domingos, Kenta Asahina, Félix Naef, Leslie B Vosshall, and Matthieu Louis. 2005. “Chemotaxis Behavior Mediated by Single Larval Olfactory Neurons in Drosophila.” Current Biology 15 (23): 2086–96. doi:10.1016/j.cub.2005.11.016.

Fishilevich, Elane, and Leslie B Vosshall. 2005. “Genetic and Functional Subdivision of the Drosophila Antennal Lobe.” Current Biology 15 (17): 1548–53. doi:10.1016/j.cub.2005.07.066.

Gao, Xiaojing J, Thomas R Clandinin, and Liqun Luo. 2015. “Extremely Sparse Olfactory Inputs Are Sufficient to Mediate Innate Aversion in Drosophila.” Edited by Matthieu Louis. PLoS ONE 10 (4). Public Library of Science: e0125986. doi:10.1371/journal.pone.0125986.

Giniger, E, S M Varnum, and M Ptashne. 1985. “Specific DNA Binding of GAL4, a Positive Regulatory Protein of Yeast.” Cell 40 (4): 767–74.

Goldman, Aaron L, Wynand van der Goes van Naters, Derek Lessing, Coral G Warr, and John R Carlson. 2005. “Coexpression of Two Functional Odor Receptors in One Neuron.” Neuron 45 (5): 661–66. doi:10.1016/j.neuron.2005.01.025.

Hallem, Elissa A, and John R Carlson. 2006. “Coding of Odors by a Receptor Repertoire.” Cell 125 (1): 143–60. doi:10.1016/j.cell.2006.01.050.

Hallem, Elissa A, Michael G Ho, and John R Carlson. 2004. “The Molecular Basis of Odor Coding in the Drosophila Antenna.” Cell 117 (7): 965–79. doi:10.1016/j.cell.2004.05.012.

Hoare, D J, C R McCrohan, and M Cobb. 2008. “Precise and Fuzzy Coding by Olfactory Sensory Neurons.” Journal of Neuroscience 28 (39): 9710–22. doi:10.1523/JNEUROSCI.1955-08.2008.

Hoare, Derek J, James Humble, Ding Jin, Niall Gilding, Rasmus Petersen, Matthew Cobb, and Catherine McCrohan. 2011. “Modeling Peripheral Olfactory Coding in Drosophila Larvae.” Edited by Bradley Steven Launikonis. PLoS ONE 6 (8): e22996–11. doi:10.1371/journal.pone.0022996.

Hodge, James J L. 2009. “Ion Channels to Inactivate Neurons in Drosophila.” Frontiers in Molecular Neuroscience 2: 1–10. doi:10.3389/neuro.02.013.2009.

Hong, Elizabeth J, and Rachel I Wilson. 2015. “Simultaneous Encoding of Odors by Channels with Diverse Sensitivity to Inhibition.” Neuron 85 (3). Elsevier Inc.: 573–89. doi:10.1016/j.neuron.2014.12.040.

Invitrogen. 2012a. “pBAD/Thio His TOPO Manual,” March, 1–74.

Invitrogen. 2012b. “pENTR™ Directional TOPO^®^ Cloning Kits,” March, 1–52.

Johns, D C, R Marx, R E Mains, B O’Rourke, and E Marbán. 1999. “Inducible Genetic Suppression of Neuronal Excitability.” Journal of Neuroscience 19 (5): 1691–97.

Jones, P L, G M Pask, and D C Rinker. 2011. “Functional Agonism of Insect Odorant Receptor Ion Channels.” In. doi:10.1073/pnas.1102425108/-/DCSupplemental/pnas.201102425SI.pdf.

Jones, Walton D, Pelin Cayirlioglu, Ilona Grunwald Kadow, and Leslie B Vosshall. 2007. “Two Chemosensory Receptors Together Mediate Carbon Dioxide Detection in Drosophila.” Nature 445 (7123). Nature Publishing Group: 86–90. doi:10.1038/nature05466.

Jung, Sarah Nicola, alexander borst, and Juergen Haag. 2011. “Flight Activity Alters Velocity Tuning of Fly Motion-Sensitive Neurons.” The Journal of Neuroscience : the Official Journal of the Society for Neuroscience 31 (25). Society for Neuroscience: 9231–37. doi:10.1523/JNEUROSCI.1138-11.2011.

Kazama, H, E Yaksi, and R I Wilson. 2011. “Cell Death Triggers Olfactory Circuit Plasticity via Glial Signaling in Drosophila.” Journal of Neuroscience 31 (21): 7619–30. doi:10.1523/JNEUROSCI.5984-10.2011.

Kazama, Hokto, and Rachel I Wilson. 2009. “Origins of Correlated Activity in an Olfactory Circuit.” Nature Neuroscience 12 (9): 1136–44. doi:10.1038/nn.2376.

Kitamoto, T. 2001. “Conditional Modification of Behavior in Drosophila by Targeted Expression of a Temperature-Sensitive Shibire Allele in Defined Neurons.” Journal of Neurobiology 47 (2): 81–92.

kitamoto, Toshihiro. 2002. “Conditional Disruption of Synaptic Transmission Induces Male-Male Courtship Behavior in Drosophila.” Proceedings of the National Academy of Sciences 99 (20): 13232–37. doi:10.1073/pnas.202489099.

Kreher, Scott A, Dennis Mathew, Junhyong Kim, and John R Carlson. 2008. “Translation of Sensory Input Into Behavioral Output via an Olfactory System.” Neuron 59 (1): 110–24. doi:10.1016/j.neuron.2008.06.010.

Krieger, J, O Klink, C Mohl, K Raming, and H Breer. 2003. “A Candidate Olfactory Receptor Subtype Highly Conserved Across Different Insect Orders.” Journal of Comparative Physiology A 189 (7): 519–26. doi:10.1007/s00359-003-0427-x.

Kwon, Jae Young, Anupama Dahanukar, Linnea A Weiss, and John R Carlson. 2007. “The Molecular Basis of CO2 Reception in Drosophila.” Proceedings of the National Academy of Sciences 104 (9): 3574–78. doi:10.1073/pnas.0700079104.

Larsson, Mattias C, Ana I Domingos, Walton D Jones, M Eugenia Chiappe, Hubert Amrein, and Leslie B Vosshall. 2004. “Or83b Encodes a Broadly Expressed Odorant Receptor Essential for Drosophila Olfaction.” Neuron 43 (5): 703–14. doi:10.1016/j.neuron.2004.08.019.

Lin, Chun-Chieh, and Christopher J Potter. 2015. “Re-Classification of Drosophila Melanogaster Trichoid and Intermediate Sensilla Using Fluorescence-Guided Single Sensillum Recording.” Edited by Matthieu Louis. PLoS ONE 10 (10). Public Library of Science: e0139675. doi:10.1371/journal.pone.0139675.

Ma, Jun, and Mark Ptashne. 1987. “The Carboxy-Terminal 30 Amino Acids of GAL4 Are Recognized by GAL80.” Cell 50 (1): 137–42. doi:10.1016/0092-8674(87)90670-2.

Nakagawa, Takao, and Leslie B Vosshall. 2009. “Controversy and Consensus: Noncanonical Signaling Mechanisms in the Insect Olfactory System.” Current Opinion in Neurobiology 19 (3): 284–92. doi:10.1016/j.conb.2009.07.015.

Neuhaus, Eva M, Günter Gisselmann, Weiyi Zhang, Ruth Dooley, Klemens Störtkuhl, and Hanns Hatt. 2005. “Odorant Receptor Heterodimerization in the Olfactory System of Drosophila Melanogaster.” Nature Neuroscience 8 (1): 15–17. doi:10.1038/nn1371.

Ng, Minna, Robert D Roorda, Susana Q Lima, Boris V Zemelman, Patrick Morcillo, and Gero Miesenböck. 2002. “Transmission of Olfactory Information Between Three Populations of Neurons in the Antennal Lobe of the Fly.” Neuron 36 (3): 463–74.

Nichols, Andrew S, and Charles W Luetje. 2010. “Transmembrane Segment 3 of Drosophila Melanogaster Odorant Receptor Subunit 85b Contributes to Ligand-Receptor Interactions.” The Journal of Biological Chemistry 285 (16): 11854–62. doi:10.1074/jbc.M109.058321.

Olsen, Shawn R, Vikas Bhandawat, and Rachel I Wilson. 2007. “Excitatory Interactions Between Olfactory Processing Channels in the Drosophila Antennal Lobe.” Neuron 54 (1): 89–103. doi:10.1016/j.neuron.2007.03.010.

Pfeiffer, B D, A Jenett, A S Hammonds, T T B Ngo, S Misra, C Murphy, A Scully, et al. 2008. “Tools for Neuroanatomy and Neurogenetics in Drosophila.” Proceedings of the National Academy of Sciences 105 (28): 9715–20. doi:10.1073/pnas.0803697105.

Pfeiffer, B D, T T B Ngo, K L Hibbard, C Murphy, A Jenett, J W Truman, and G M Rubin. 2010. “Refinement of Tools for Targeted Gene Expression in Drosophila.” Genetics 186 (2): 735–55. doi:10.1534/genetics.110.119917.

Pulver, Stefan R, Stanislav L Pashkovski, Nicholas J Hornstein, Paul A Garrity, and Leslie C Griffith. 2009. “Temporal Dynamics of Neuronal Activation by Channelrhodopsin-2 and TRPA1 Determine Behavioral Output in Drosophila Larvae.” Journal of Neurophysiology 101 (6). American Physiological Society: 3075–88. doi:10.1152/jn.00071.2009.

Robertson, Hugh M, Coral G Warr, and John R Carlson. 2003. “Molecular Evolution of the Insect Chemoreceptor Gene Superfamily in Drosophila Melanogaster.” Proceedings of the National Academy of Sciences 100 Suppl 2 (Supplement 2): 14537–42. doi:10.1073/pnas.2335847100.

Silbering, A F, R Rytz, Y Grosjean, L Abuin, P Ramdya, G S X E Jefferis, and R Benton. 2011. “Complementary Function and Integrated Wiring of the Evolutionarily Distinct Drosophila Olfactory Subsystems.” Journal of Neuroscience 31 (38): 13357–75. doi:10.1523/JNEUROSCI.2360-11.2011.

Smart, Renee, Aidan Kiely, Morgan Beale, Ernesto Vargas, Colm Carraher, Andrew V Kralicek, David L Christie, Chen Chen, Richard D Newcomb, and Coral G Warr. 2008. “Drosophila Odorant Receptors Are Novel Seven Transmembrane Domain Proteins That Can Signal Independently of Heterotrimeric G Proteins.” Insect Biochemistry and Molecular Biology 38 (8): 770–80. doi:10.1016/j.ibmb.2008.05.002.

Song, Z, and H Steller. 1999. “Death by Design: Mechanism and Control of Apoptosis.” Trends in Cell Biology 9 (12): M49–M52.

Stocker, R F, M C Lienhard, A Borst, and K F Fischbach. 1990. “Neuronal Architecture of the Antennal Lobe in Drosophila Melanogaster.” Cell and Tissue Research 262 (1). Springer-Verlag: 9–34. doi:10.1007/BF00327741.

Su, Chih-Ying, Karen Menuz, Johannes Reisert, and John R Carlson. 2012. “Non-Synaptic Inhibition Between Grouped Neurons in an Olfactory Circuit.” Nature 492 (7427). Nature Publishing Group: 66–71. doi:10.1038/nature11712.

Sweeney, Sean T, Kendal Broadie, John Keane, Heiner Niemann, and Cahir J O’Kane. 1995. “Targeted Expression of Tetanus Toxin Light Chain in Drosophila Specifically Eliminates Synaptic Transmission and Causes Behavioral Defects.” Neuron 14 (2): 341–51. doi:10.1016/0896-6273(95)90290-2.

van der Bliek, A M, and E M Meyerowitz. 1991. “Dynamin-Like Protein Encoded by the Drosophila Shibire Gene Associated with Vesicular Traffic.” Nature 351 (6325): 411–14. doi:10.1038/351411a0.

Vosshall, L B, A M Wong, and R Axel. 2000. “An Olfactory Sensory Map in the Fly Brain.” Cell 102 (2): 147–59.

Vosshall, L B, and B S Hansson. 2011. “A Unified Nomenclature System for the Insect Olfactory Coreceptor.” Chemical Senses 36 (6): 497–98. doi:10.1093/chemse/bjr022.

Vosshall, Leslie B, Hubert Amrein, Pavel S Morozov, Andrey Rzhetsky, and Richard Axel. 1999. “A Spatial Map of Olfactory Receptor Expression in the Drosophila Antenna.” Cell 96 (5): 725–36. doi:10.1016/S0092-8674(00)80582-6.

Wagh, Dhananjay A, Tobias M Rasse, Esther Asan, Alois Hofbauer, Isabell Schwenkert, Heike Dürrbeck, Sigrid Buchner, et al. 2006. “Bruchpilot, a Protein with Homology to ELKS/CAST, Is Required for Structural Integrity and Function of Synaptic Active Zones in Drosophila.” Neuron 49 (6): 833–44. doi:10.1016/j.neuron.2006.02.008.

Wilson, Rachel I. 2011. “Understanding the Functional Consequences of Synaptic Specialization: Insight From the Drosophila Antennal Lobe.” Current Opinion in Neurobiology 21 (2). Elsevier Ltd: 254–60. doi:10.1016/j.conb.2011.03.002.

Yaksi, Emre, and Rachel I Wilson. 2010. “Electrical Coupling Between Olfactory Glomeruli.” Neuron 67 (6). Elsevier Inc.: 1034–47. doi:10.1016/j.neuron.2010.08.041.

